# “Evaluation of cytotoxic effects of diverse essential oils on human keratinocytes (HaCaT)”

**DOI:** 10.1101/2025.04.16.649163

**Authors:** Alicja Karolina Surowiak, Marcin Poręba, Daniel Jan Strub

## Abstract

Essential oils (EOs) are widely used in cosmetics for their active and aromatic properties, yet their cytotoxicity remains a concern, especially regarding skin safety. This study aimed to evaluate the cytotoxicity of 32 essential oils on immortalized keratinocytes (HaCaT cells) using an MTS assay. Results demonstrated significant variability in cell viability, with 14 oils reducing viability below 50%. These findings underscore the need for further research into the specific chemical constituents responsible for the observed effects.

## Introduction

Natural complex substances (NCSs), including essential oils (EOs), are extensively utilized across various sectors such as food, cosmetics, and pharmaceuticals due to their diverse biological properties. The cosmetics industry warrants special attention, as nature-based products are a growing trend. The natural cosmetics market was valued at 45.2 billion USD in 2023, with projections for growth to 76.5 billion USD by 2033, at a compound annual growth rate (CAGR) of 5.4% [1]. Moreover, one-third of new cosmetic products launched in EU in 2024 had eco-certifications [2]Which emphasize the need for research on natural products. EOs are widely incorporated into cosmetics as active ingredients, fragrances, and aromatherapeutic agents in both skincare and haircare products. The skin, as the primary interface between NCSs and the human body (following the respiratory system), is the largest organ and a crucial barrier against external factors. Direct contact between the skin and NCSs can lead to various pathological conditions, including allergies and eczema. To avoid these the use of EOs is regulated by organizations such as the International Fragrance Association (IFRA), which sets guidelines for the safe concentration levels of various EOs in cosmetic formulations. Additionally, EU and FDA regulations apply, nevertheless this do not eliminate the risk of side effects. Given the widespread incorporation of NCSs, particularly EOs, into consumer products, it is essential to assess their safety profile.

The aim of this study was to evaluate the cytotoxicity of 32 essential oils, focusing on their ability to reduce the viability of immortalized keratinocytes (HaCaT cells). This research forms the foundation for further investigation into the safety and biological activity of EOs, providing a more accurate understanding of their potential effects on human skin.

## Materials

## Cell line and reagents

This study was carried out using HaCaT cell lines from DKFZ Heidelberg (CLS Cell Lines Service GmbH, d/b/a Cytion). HaCaT cells were cultured in Dulbecco’s Modified Eagle’s Medium (DMEM) (CLS Cell Lines Service GmbH, d/b/a Cytion) w: 4.5 g/L Glucose, w: 4 mM L-Glutamine, w: 1.5 g/L NaHCO3, w: 1.0 mM Sodium pyruvate supplemented with 10% Fetal Bovine Serum (heat inactivated) (FBS, Gibco Life Technologies), 100units/mL penicillin and 10 ug/mL streptomycin (both Gibco Life Technologies) in a humidified 5% CO2 atmosphere at 37 °C. MTS was obtained from Promega (CellTiter 96® AQueous One Solution Cell Proliferation Assay). Dimethyl sulfoxide (DMSO) was purchased from Honeywell Sp. z.o.o.

Thirty-two samples of natural fragrance materials were generously donated by the essential oil industry, members of the International Federation of Essential Oils and Aroma Trades, to ensure the highest quality. These materials included:

**Table 1.** List of the NCSs used in the study.

| Name | Company | CAS Number | Botanical name | Origin |
| --- | --- | --- | --- | --- |
| Pinus ponderosa EO | Hierbas Patagónicas SRL | 97553-47-4 | *Pinus ponderosa* | Argentina |
| Petitgrain lemon EO | Bordas S.A. | 8048-51-9 | *Citrus limon (L.) Osbeck* | Spain |
| Clove Leaf EO | Jacaradnas International | 8000-34-8 | *Eugenia caryophyllata L.* | Madagascar |
| Hiba EO | Bristol Botanicals Ltd. | 68917-43-1 | *Thujopsis dolabrata (Thunb. ex L. f.) Siebold & Zucc.* | Japan |
| Tea tree EO | The Lebermuth Co. | 68647-73-4 | *Melaleuca alternifolia (Maiden & Betche) Cheel* | China |
| Sandalwood EO | Kallin International Ltd. | 8024-35-9 | *Santalum album L.* | India |
| Ylang Ylang I EO | Jacaradnas International | 83863-30-3/<br>8006-81-3 | *Cananga odorata Hook. f. & Thomson forma genuina* | Madagascar |
| Lemongrass EO | A. Fakhry & Co. | 8007-02-01 | *Cymbopogon flexuosus (Nees ex Steud.) W. Watson* | Egypt |
| Lavender EO | EO of Tasmania | 8000-28-0 | *Lavandula angustifolia* | Australia |
| Australian Blue Cypress EO | Essentially Australia | 187348-13-6 | *Callitris intratropica Baker & H.G.Sm.* | Australia |
| Patchouli Oil Sumatra Iron Free Min 32 PA EO | PT Van Aroma | 84238-39-1 / 8014-09-3 | *Pogostemon cablin (Blanco) Benth.* | Indonesia |
| Eucalyptus globulus EO | EO of Tasmania | 84625-32-1 | *Eucalyptus globulus* | Australia |
| Cedarwood Himalayan EO | The Lebermuth Co. | 68991-36-6 | *Cedrus deodora (Roxb. ex Lamb.) G.Don* | India |
| Ravintsara EO | Jacaradnas International | 92201-50-8 | *Cinnamomum camphora Ness et Eberm.* | Madagascar |
| White Oud EO | Hermitage Oils SRL | 1333524-00-7 | *Aetoxylon Sympetalum* | Indonesia |
| Clove Bud EO | Jacaradnas International | 8000-34-8 | *Syzygium aromaticum (L.) Merr. & L. M. Perry syn. Eugenia caryophyllus (Spreng.) Bullock & S. G. Harrison* | Madagascar |
| Calendula EO | A. Fakhry & Co. | 84776-23-8 | *Calendula officinalis L.* | Egypt |
| Thyme EO | Vessel Essential Oils | 84929-51-1 | *Thymus vulgaris L.* | Greece |
| Siam wood EO | Berje Inc. | 8022-47-7 | *Fokienia hodginsii (Dunn) A.Henry & H. H.Thomas* | Vietnam |
| Gurjun Dark – Gurjunene EO | PT Van Aroma | 8030-55-5 | *Dipterocarpus C. F. Gaertner* | Indonesia |
| Buddha Wood EO | Essentially Australia | 1429902-59-9 | *Eremophila mitchellii Benth.* | Australia |
| Rosewood (Bois de rose) EO | Berje Inc. | 8015-77-8 | *Aniba rosodora Ducke* | India |
| Cedarwood Heart EO | Hermitage Oils SRL | 8023-85-6 | *Cedrus atlantica* | Morocco |
| Jasmine EO | A. Fakhry & Co. | 84776-64-7 / 8022-96-6 | *Jasminum grandiflorum L.* | Egypt |
| Turmeric EO | Bristol Botanicals Ltd. | 8024-37-1 | *Curcuma longa L.* | Unknown |
| Cedrat EO | Hermitage Oils SRL | 68991-25-3 | *Citrus medica* | Sicily |
| Wild carrot seed EO | Vessel Essential Oils | 8015-88-1 | *Daucus carota subsp. sativus* | Macedonia |
| Kaffir Lime Leaf EO | PT Van Aroma | 91771-50-5 | *Citrus hystrix* | Indonesia |
| Savory summer EO | Berje Inc. | 8016-68-0 | *Satureja hortensis L.* | Hungary |
| Rose centifolia EO | A. Fakhry & Co. | 8007-01-0 | *Rosa x centifolia L. syn. Rosa gallica var. centifolia L.* | Egypt |
| Peppermint EO | A. Fakhry & Co. | 8006-90-4 | *Mentha x piperita L.* | Egypt |
| Opoponax EO extra | Payan Bertrand S.A. | 8021-36-1 | *Commiphora erythraea var. glabrescens Engler* | Somalia |

## Methods

EOs were diluted in DMSO to initial concentration of 10 mg/mL.

Cell viability MTS assay

HaCaT cells were seeded in 96-well cell culture plates (50,000 cells per well in 100 μL of phenol red-free DMEM medium) and allowed to attach overnight. The next day, the medium was changed and the cells were treated with test substances (200 μg/mL) for 24 h. After this time, cell viability was determined using an MTS assay according to the manufacturer’s protocol (CellTiter 96® Aqueous One Solution Cell Proliferation Assay). The number of living cells is reported as a survival ratio calculated based on the absorbance value measured in control wells (untreated cells). All measurements were performed in triplicate.

## Results and discussion

Of the thirty-two EOs tested, fourteen caused a reduction in the viability of immortalized keratinocytes (HaCaT) to below 50%. The cytotoxicity of EOs results from a complex interaction between their chemical composition, plant origin, and metabolic processes. Both the presence and concentration of specific compounds significantly impact the biological activity and safety of these oils.

**Table 2.** Results of cell viability against HaCaT, concentration 200 μg/mL time of exposition 24h.

| Name of the NCS | Cell viability [%] | SD |
| --- | --- | --- |
| Pinus ponderosa EO | 83.04 | 6.66 |
| Petitgrain lemon EO | 30.67 | 3.30 |
| Clove Leaf EO | 84.07 | 2.75 |
| Hiba EO | 24.94 | 3.96 |
| Tea tree EO | 90.95 | 4.46 |
| Sandalwood EO | 23.16 | 0.25 |
| Ylang Ylang I EO | 24.97 | 1.80 |
| Lemongrass EO | 43.48 | 1.82 |
| Lavender EO | 88.68 | 2.08 |
| Australian Blue Cypress EO | 25.80 | 3.08 |
| Patchouli Oil Sumatra Iron Free Min 32 PA EO | 21.15 | 4.58 |
| Eucalyptus globulus EO | 90.96 | 4.19 |
| Cedarwood Himalayan EO | 32.70 | 2.95 |
| Ravintsara EO | 94.33 | 2.18 |
| White Oud EO | 23.32 | 2.43 |
| Clove Bud EO | 93.45 | 3.74 |
| Calendula EO | 23.38 | 2.21 |
| Thyme EO | 78.06 | 1.16 |
| Siam wood EO | 78.64 | 5.01 |
| Gurjun Dark – Gurjunene EO | 21.61 | 3.33 |
| Buddha Wood EO | 29.37 | 0.86 |
| Rosewood (Bois de rose) EO | 70.95 | 1.26 |
| Cedarwood Heart EO | 22.50 | 0.52 |
| Jasmine EO | 91.64 | 5.80 |
| Turmeric EO | 90.81 | 3.02 |
| Cedrat EO | 96.69 | 2.44 |
| Wild carrot seed EO | 92.34 | 2.01 |
| Kaffir Lime Leaf EO | 75.51 | 5.13 |
| Savory summer EO | 99.41 | 6.02 |
| Rose centifolia EO | 92.52 | 1.97 |
| Peppermint EO | 95.90 | 5.07 |
| Opoponax EO extra | 24.64 | 3.40 |

Among the most toxic essential oils, those rich in sesquiterpenes (Fig. 1) and phenols can be identified. Sandalwood essential oils previously showed high cytotoxicity (0.30%), resulting in approximately 10% cell viability towards HaCaT cells [3], which corresponds with our results.

**Figure 1.**
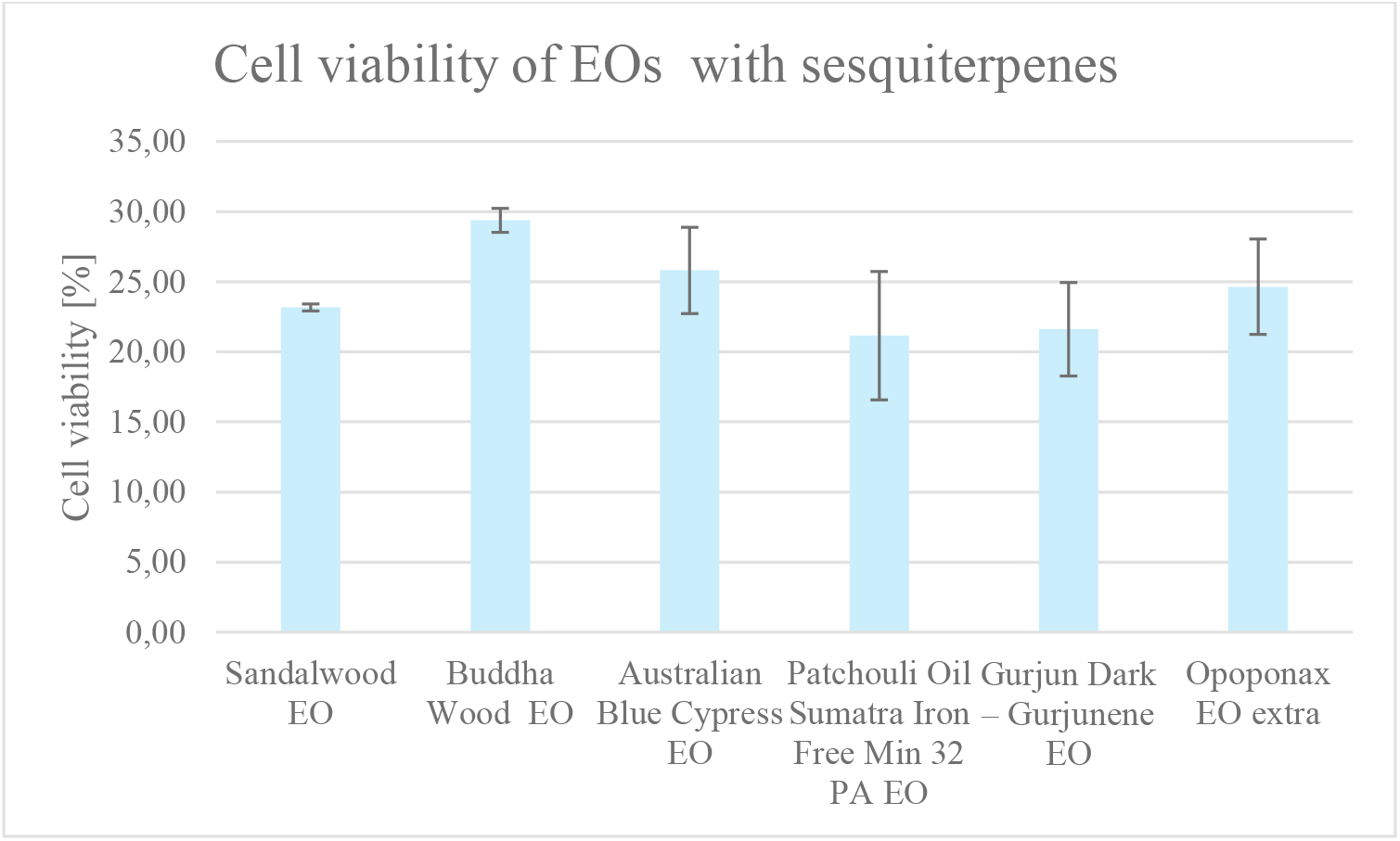
HaCaT cell viability at 200 µg/mL after 24h exposure of essential oil rich in sesquiterpenes.

Among the cytotoxic EOs, we identified Ylang-Ylang EO, whose main constituents are monoterpenes. Previously, Karabatak et al. reported a cell viability of approximately 55.75% at a concentration of 250 µg/mL [4], which is similar to the concentration tested in this study, although our cell viability was more than twice as low. The difference may be due to the fact that our EOs were derived from the first fraction of distilled oils, whereas the authors did not specify whether the commercial EO was a fraction or the whole oil. This could explain the significant difference in chemical composition. Among the EOs with minimal effects on the viability of immortalized keratinocytes, only a few have been previously tested. Tea tree EO showed no toxicity to cells (∼90% cell viability) in earlier studies. However, in one study, a much lower concentration (0.00–0.25% w/v) was analyzed [5], while in another study, a concentration half as low as in our experiment (100 µg/mL) also showed no effects[6]. Moreover, the IC_50_ value in previous studies was found to be more than four times higher than in our experiment (930 μg/mL) [7]. The essential oil of *Lavandula angustifolia* was previously analyzed on the HaCaT cell line. In our study, a 24-hour exposure to 200 μg/mL resulted in approximately 88% cell viability, whereas previous studies reported only 25% cell viability at a concentration of 1% [8], with an LC_50_ value ranging from 0.95 to 1.48 μL/mL [9] and IC_50_ values of 0.43% [5] and 0.2% [10] in other studies. A similar situation was observed with thyme and peppermint EOs. The IC_50_ value for thyme EO was reported as 0.18% [5], and the LC_50_ value of peppermint EO was 2.02 μL/mL [9], while in our study, cell viability was approximately 78% for thyme EO and 96% for peppermint EO. Turmeric essential oil showed no impact on cell viability at a concentration of 1% in a previous study [11] However, in another study, the reported IC_50_ value (56.1 µg/mL) [12] was almost four times higher than in our study, which resulted in ∼90% cell viability. In the same study, ravintsara EO was analyzed, with an estimated IC_50_ value of 250.90 μg/mL [12]. The previously reported cytotoxicity of *Eugenia caryophyllata* L. essential oils has primarily focused on clove buds. The reported IC_50_ value was estimated at 0.0099% [5], while in other studies, a concentration of 0.5% had no impact on cell viability [13], and 0.625% resulted in ∼7% cell viability [14]. There are no available literature data on the cytotoxicity of clove leaf essential oil. In our study, clove bud and clove leaf EOs showed 93% and 84% cell viability, respectively.

Interesting results were obtained for EOs containing citral and limonene, substances known for causing side effects while applied in cosmetic products. Among the tested EOs, citral is the dominant component of lemongrass EO[15], while limonene is the main constituent of petitgrain lemon [16] and kaffir lime leaf EOs [17]. Cedrat EO typically contains both ingredients in variable proportions [18]. In our study, only petitgrain lemon essential oil exhibited significant cytotoxicity (Fig. 2), which corresponds with previously reported results [16].

**Figure 2.**
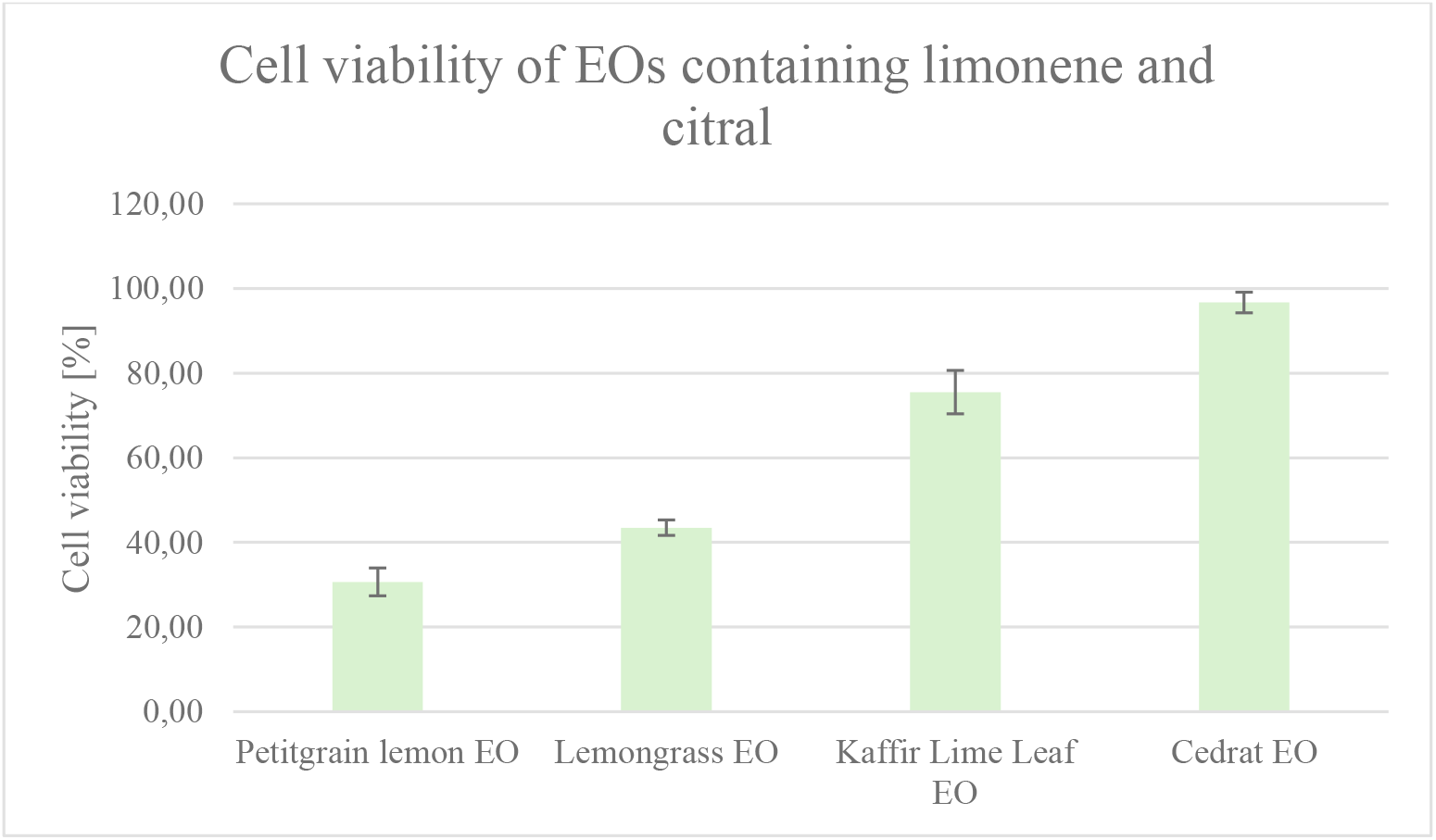
Comparison of cell viability of EOs rich in limonene and citral.

IC_50_ values for lemongrass essential oil (EO) vary. Our research demonstrated 43.48% cell viability at a concentration of 200 μg/mL. Al-Ghanayem et al. estimated the IC_50_ at 1250 μg/mL [19], whereas Kozics et al. reported a value of 0.0056% [5], both using the MTT assay. Cedrat EO had almost no impact on cell viability in our study, whereas previously reported IC_50_ values were nearly 10 times lower (∼24 μg/mL) [20] than the concentration used in our study. The differences in the effects of EOs on immortalized keratinocytes may result from multiple factors. Primarily, they depend on the raw material itself—its geographical origin, harvesting method, the plant part used, and the extraction process. These factors influence the chemical composition of EOs, affecting both the synergistic and antagonistic interactions between their components. Additionally, slight variations in experimental conditions, such as cell culture parameters, exposure time, and assay type, may impact the results. Another significant source of variation arises from differences between researcher-obtained samples and commercial essential oils. Manufacturers often standardize their products to ensure consistency across batches, whereas smaller producers may not follow such practices. Furthermore, commercial oils may contain additives from uncertain sources. In contrast, experimental EOs are non-standardized samples obtained under specific conditions, which may further contribute to the observed differences in their biological effects. The exact relationships between the components of individual essential oils will be analyzed following composition analysis.

## Conclusion

This study underscores the complexity of evaluating the safety of natural complex substances like EOs, which are widely used in the cosmetic industry. Further research is needed to explore the underlying mechanisms and the role of specific compounds in modulating the cytotoxic effects of EOs. This research will be crucial in guiding the safe incorporation of EOs into cosmetic products and ensuring their regulatory compliance.

## Funding

This research received no external funding.

## Data Availability Statement

Not applicable.

## Acknowledgments

We would like to thank representatives of the essential oils industry and members of the International Federation of Essential Oils and Aroma Trades (IFEAT): A. Fakhry & Co., Berje Inc., Bordas S.A., Bristol Botanicals Ltd., EO of Tasmania, Essentially Australia, Hermitage Oils SRL, Hierbas Patagónicas SRL, Jacaradnas International, Kallin International Ltd., Payan Bertrand S.A., PT Van Aroma, The Lebermuth Co., Vessel Essential Oils.

## Conflicts of Interest

The authors declare no conflict of interest.

## Notes

### Competing Interest Statement

The authors have declared no competing interest.

